# shRNAI: a deep neural network for the design of highly potent shRNAs

**DOI:** 10.1101/2024.01.09.574789

**Authors:** Seokju Park, Sung-Ho Park, Jin-Seon Oh, Yung-Kyun Noh, Junho K Hur, Jin-Wu Nam

## Abstract

miRNA-mimicking short hairpin RNA (shRNAmir), which depends on whole miRNA biogenesis, has been used to elucidate the function of target genes and to develop therapeutic approaches due to its stable, robust downregulation. Despite efforts to design potent shRNAmir guide RNAs (gRNAs), biological features other than the sequence have not been fully considered. Here, we developed shRNAI, a convolutional neural network model for the prediction of highly potent shRNAmir gRNAs. The shRNAI model trained with gRNA sequences alone predicted more potent shRNAmir gRNAs than the existing algorithms. The shRNAI+ model, trained with the additional features of shRNAmir processibility and target site context, was further improved on both public and our experimental test datasets. Although the shRNAI models were trained on datasets generated with the CNNC motif-free shRNAmir backbone, they also exhibited better performance for the CNNC motif. Our study provides not only a rational framework for designing shRNAmir gRNAs for targets but also a means of designing optimal RNA interference drugs necessary for RNA therapeutics.

## INTRODUCTION

RNA interference (RNAi) has been widely used to downregulate target genes in human cells since sequence-specific gene silencing was reported in animals (1,2). Short interfering RNAs (siRNAs), short hairpin RNAs (shRNAs) (3), and miRNA-mimicking short hairpin RNAs (shRNAmirs) (4) have been developed as RNAi platforms with respective advantages and disadvantages. Similar to primary miRNA (pri-miRNA), shRNAmir is processed at a specific position in the basal stem by Drosha in the nucleus (5–9), followed by processing in the apical stem by Dicer in the cytoplasm (10–22). A guide RNA (gRNA) derived from the remaining duplex is loaded onto Argonaute (AGO) to silence its target, which is a similar process to that of miRNA (23–29). Although miRNA targeting requires only partial base pairing (at positions 2-7 from the 5’-end of the miRNA) (30–34), RNAi requires nearly complete base pairing to induce the endonucleolytic cleavage ability of AGO (35,36).

While siRNA is manufactured in the form of duplex RNA and injected into cells, shRNAmir is expressed through a DNA vector and processed via the endogenous miRNA pathway. Due to its vector-based stable expression and ability to utilize the endogenous processing pathway, shRNAmir has been known to exhibit greater potency in repressing target genes than other platforms both *in vitro* and *in vivo* (37–41). Since Zeng *et al* first succeeded in modeling an artificial pri-miR-30-based shRNAmir and selectively repressing the target (4), multiple attempts have been made to design more potent shRNAmirs by engineering pri-miRNAs, including pri-miR-30 backbones (42–53). In terms of both Drosha processing efficiency and accuracy, pri-miR-30 and its engineered backbones seemed to be superior to the other miRNAs (54). However, because the first pri-miR-30 backbone was mutated to insert an EcoR1 restriction enzyme site in the CNNC motif (55) (Supplementary Figure 1A), which is an important cis-acting element in Drosha processing, the restriction enzyme site was moved downstream in a subsequent version, yielding the miR-E backbone (56).

In addition to backbone engineering, algorithms for optimizing gRNA sequences have been developed to design potent shRNAmirs. Previously developed prediction tools for the potency of gRNAs using mir-30/mir-E backbones utilized shallow machine learning algorithms with k-mer-based features of RNA (54,57,58). However, due to the restricted backbones, the sequence and structure of the respective gRNA:passenger duplex can impact the processing and targeting of shRNAmirs, indicating that the optimal gRNA:passenger duplex for a given backbone must be carefully selected among candidates.

In this study, we developed shRNAI, which implements a convolutional neural network (CNN) to accurately predict the potency of gRNAs embedded in the pri-miR-30 backbone. Large shRNAmir datasets produced through massive parallel assays were precompiled. The original shRNAI model outperformed existing algorithms, but this model was improved by adding more features, including processing efficiency and site context information and *in vitro* and *in silico* benchmarking. The improved shRNAI+ model is available at http://big2.hanyang.ac.kr/shRNAI.

## MATERIALS AND METHODS

### Public data sources

The gene annotation data were downloaded from the GENCODE database (https://www.gencodegenes.org/) using GENCODE v41 and vM30 for the development of the shRNAI models and for predicting shRNAI scores for all human and mouse genes. *In vitro* Drosha processing data were downloaded from SRA051323 of the Short Read Archive (https://www.ncbi.nlm.nih.gov/sra/).

### Training and evaluation datasets

For training and testing of the shRNAI models, shRNA potency and shRNAmir processing data were collected from previous papers (55,58–60) and the National Center for Biotechnology Information (NCBI) Gene Expression Omnibus (http://www.ncbi.nlm.nih.gov/geo/; GSE62183). Approximately 300k combined data points were precompiled from the shERWOOD (S), TILE (T), M1 (M), Ras (R), miR-E (E), and UltramiR (U) datasets. The S, T, and M datasets were selected for training and hold-out tests because they have more data to train a deep learning model, and the other datasets were used as independent test data. Before training the models, all redundant data points among the datasets were removed, and min–max scaling was applied to each dataset. For evaluation, shRNA screening datasets were collected from previous studies (61–65).

### Building shRNAI models

#### Architecture

The shRNAI models were based on a CNN with three convolutional layers (66–68). The input layers received a four-by-*n* matrix through one-hot encoding, which was transformed from the input gRNA sequence, with a length of 22 base pairs (bp), or from its target site, with a length of 22 bp + a flanking region of 14 bp. The number of convolutional layers was tuned by considering the number of parameters in the networks and the cross-validation results. The batch-normalized output of each convolution layer was activated through an exponential linear unit (ELU) and input into the next layer (69,70). The number of convolution filters was chosen to be [128, 256]. The column size of the filters in each convolution layer was set as [3, 5, 7], and the row size was set to 1, except for the first convolution layer, which was set to 4 to cover all 4 nucleotide inputs. Between the first and second convolution layers, we adopted a max pooling layer with a filter of size 2 and a stride of 2. The average pooling result of the final convolution layer output was input into two fully connected layers of [64, 128, 256] hidden ELU units with batch normalization. For the shRNAI+ model, additional features beyond the sequence input were concatenated with the output of the final convolution layer. The final regression output layer performed a linear transformation of the previous layer’s output to predict the potency.

#### Training

The network was trained by the Adam optimizer to minimize the mean-square error relative to the observed values in the training data. We used the Keras Python library with the TensorFlow backend to implement the CNN (71,72). The training continued for up to 50 epochs but was stopped early if there was no loss reduction in the validation dataset for at least 6 consecutive epochs. The initial learning rate in Adam was set to 0.001 and decayed by a factor of 0.1 after no improvement for 3 epochs.

We set the hyperparameters, including the convolution filter width, number of convolution filters, and number of fully connected layer neurons, by 5-fold cross-validation on 90% of the training dataset. The other 10% of the training dataset was used as the hold-out test dataset to evaluate the final model.

### Prediction model for Drosha substrate surplus

#### Architecture

The prediction model for Drosha substrate surplus was also based on a CNN. The input was constructed by concatenating two consecutive sequences. One sequence ranged from 27 nucleotides (nt) upstream to 29 nt downstream of the 5’ Drosha cleavage site, and the other sequence ranged from 31 nt upstream to 25 nt downstream of the 3’ Drosha cleavage site. Then, the reverse sequence of the latter was added to the former sequence. After transforming it by one-hot encoding, an eight-by-*n* matrix was constructed of the length of the input features. The final input was constructed by adding the base-pair information, resulting in a nine- by-*n* matrix. The CNN was constructed with the three convolutional layers that achieved the best performance on the validation dataset. Batch normalization was adopted between convolution and rectified linear unit (RELU) (73) activation in each convolutional layer. The number of convolution filters was chosen to be [128, 256, 512]. The column size of the filters in each convolution layer was chosen to be [(3,1), (5,1), (7,1)] except for the first convolution layer, which was set to (9,9) to cover all nine input dimensions. Between the convolution layers, we adopted a max pooling layer with a filter of size 2 and a stride of 2. The average pooling result of the final convolution layer was input into two fully connected layers of [64, 128] hidden RELU units via batch normalization. The final regression output layer performed a linear transformation of the output of the previous layer to predict the Drosha substrate surplus.

#### Training

The CNN model was trained as described for the shRNAI model above.

### Testing shRNAI on screening data

Before testing the models on the screening data, we collected only the shRNAs in which the target sites were present in the gene annotation data. When the flanking sequence of the target site was less than 14 nt in length, the remaining sequence was filled with random nucleotides.

#### Large-scale screening dataset

The phenotypes of each shRNA and each gene were calculated as previously published (58), with some exceptions. The phenotypes for each shRNA were calculated as the mean log2-fold change for the two replicates. To test the retrospective prediction power of the algorithms, the observed phenotypes of the top and bottom predictions were compared using only previously annotated essential genes. To test the prediction accuracy of the algorithms, gene-level scores that were calculated from the mean phenotypes for the top and bottom predictions were compared to identify essential or nonessential genes. The dataset included ∼25 shRNAs per candidate target gene, although there were some redundancies between the sublibraries.

#### Small-scale screening dataset

Data generated (1) with more than 100 target genes for a robust comparison and (2) with the negative selection method were collected to benchmark our model. The Rathert 2015 dataset included 2,917 mir-30- based shRNAmirs without the CNNC motif to identify chromatin factors that prevent resistance to the bromodomain in acute myeloid leukemia (62) (Supplementary Figure 3B-D). The Huang 2014 dataset included 2,245 mir-30-based shRNAmirs without the CNNC motif targeting 442 known druggable target genes in MP1 cell lines (63) (Supplementary Figure 3E-G). The Banito 2018 dataset included ∼2,400 CNNC motif-free mir-30-based shRNAmirs targeting 400 chromatin regulators to identify key regulators involved in sustaining synovial sarcoma cell transformation (64) (Supplementary Figure 3H-L). The David 2016 dataset included 558 mir-E backbone shRNAmirs targeting 93 transcription factors (65) (Supplementary Figure 3M-O). All the datasets were preprocessed via the same method. The phenotypes of each shRNA were calculated as the mean log2-fold change for all replicates after normalization to the total reads of each replicate. Gene-level scores were calculated as the mean phenotypes for all tested shRNAs of each gene. For the evaluation of the algorithms, the observed phenotypes of the top and bottom predictions were compared in the most negative subset of tested genes.

### Cell culture and lentivirus production

A human embryonic kidney 293T (HEK293T) cell line, a HeLa cell line, and a Huh7 cell line were cultured at 90% confluence in media consisting of high-glucose Dulbecco’s modified eagle medium (DMEM), 10% fetal bovine serum (FBS), 100 U/mL penicillin/streptomycin and 2.5 mg antimicrobial reagent. The cells were treated with 0.05% trypsin (Welgene, Korea) and subcultured at a 1/10 volume. HEK293T cells were seeded in 100 mm dishes (NUNC, Thermo Fisher Scientific, Denmark) at a density of 3 x 10^5 cells/well and transfected using Lipofectamine 2000 (Invitrogen, USA) according to the manufacturer’s instructions. HEK293T cells were transfected with 1.3 pmol of psPAX2 (Addgene #12260), 1.72 pmol of pMD2.G (Addgene # 12259), and 1.64 pmol of the transfer plasmid for each shRNA expression vector. The transfected cells were cultured for 72 h, after which virus preparation was performed.

### Lentiviral transduction and quantification of RNAi efficiency

The HeLa cells and Huh7 cells were seeded in 12-well plates (NUNC, Thermo Fisher Scientific, Denmark) at a density of 0.8 x 10^5 cells/well and transfected with 10 µg/ml polybrene (Sigma‒Aldrich, USA). The transfected cells were cultured for 72 h, after which puromycin (1 µg/ml) (Gendipo, Korea) selection was performed for 48 h, and the cells were transferred to 6-well plates. The transfected cells were treated with doxycycline (1 µg/ml) (Sigma‒Aldrich, USA) for 48 h, after which RNA preparation was performed. An easy-spin total RNA extraction kit (Intron, Korea) was used for RNA preparation according to the manufacturer’s instructions. cDNA was synthesized using a high-capacity cDNA reverse transcription kit (Thermo Fisher Scientific, Denmark) with 1 µg of total RNA according to the manufacturer’s instructions. The RNA expression level was analyzed via qRT‒PCR (THUNDERBIRD™ Next SYBR® qPCR Mix (TOYOBO, Japan)).

### RNA sequencing processing for off-target analysis

RNA sequencing (RNA-seq) data were processed to remove adapter sequences from reads using Cutadapt (version 2.10, parameter: -m 25) (74). Processed RNA-seq reads were mapped, and gene expression levels were quantified using RSEM with the GRCH38 genome and GENCODE v38 gene annotation (version 1.3.1, parameter: --star) (75). To compare gene expression levels, fragments per kilobase of exons per million mapped reads (FPKM) values of all genes were normalized by quantile normalization, and genes with more than 1 FPKM in both the negative control and shRNAmir-transfected samples were used for further analysis.

## RESULTS

### shRNAI pipeline for selecting potent gRNAs

To design potent gRNAs and their backbones, we developed the shRNAI pipeline comprising the following five steps: 1) selecting a target gene and region; 2) listing all possible gRNA candidates of 22 nt or 50 nt tiling the target region; 3) extracting features for the shRNAI model; 4) sorting candidates by scoring using the shRNAI model; and 5) generating an shRNAmir by inserting the best gRNA duplex into miRNA-based backbones (Figure 1A).

**Figure 1.**
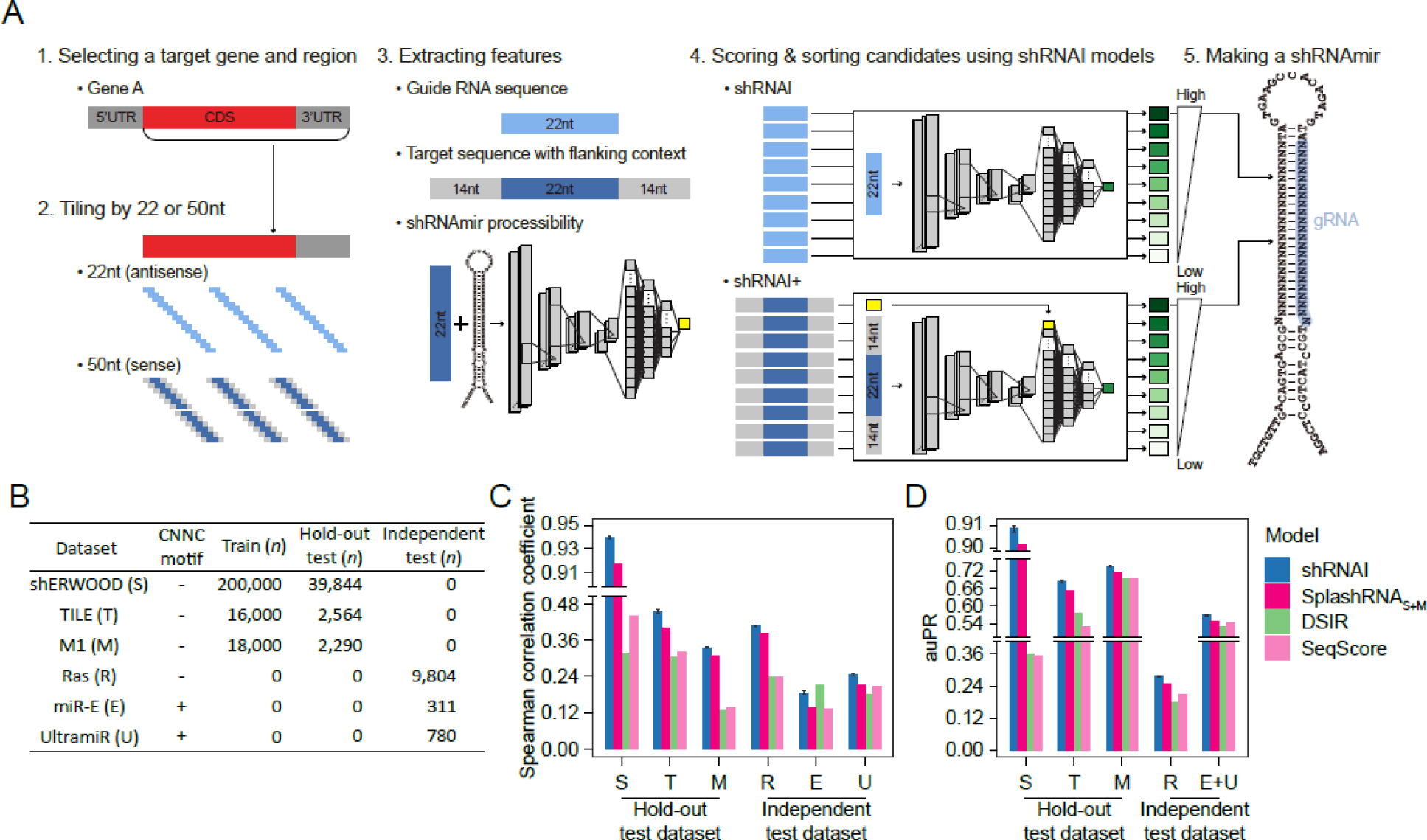
A deep learning model was constructed to predict shRNA potency. (A) A schematic workflow for the shRNAmir design process of shRNAI. (B) The number of full data points for model training and testing. To implement the model, the shERWOOD, TILE, and M1 datasets were selected and randomly divided into training and hold-out test sets. The Ras, miR-E, and UltramiR datasets were used as independent test sets. (C and D) Performance comparison of regression (C) and classification (D) of shRNAI with other algorithms using test sets. For shRNAI, the data represent the median ± SD for 100 models trained with the same hyperparameters to show the convergence of the stochastic nature of building deep learning models.

To construct an shRNAI model to predict gRNA potency, datasets from previous shRNAmir studies in which the knockdown efficiency of each pri-miR-30-based shRNAmir was measured via a massively parallel assay were collected (Figure 1B). CNN-based models were trained using balanced training datasets from the S, T, and M datasets to select the dataset that exhibited the best performance (Supplementary Figure 1B and C). The largest dataset, S, which exhibited the best performance in the hold-out test dataset, and the second largest dataset, M, which exhibited the best performance in the independent test dataset, were selected as the final training dataset for the shRNAI model (Supplementary Figure 1D and E).

shRNAI was benchmarked with the current state-of-the-art algorithms SplashRNA (58), DSIR (57), and SeqScore (54). To ensure a fair comparison, SplashRNA was retrained with the S+M dataset, named SplashRNAS+M, with the same parameters used in the original SplashRNA paper. shRNAI outperformed the other models for both the mir-30 and mir-E datasets according to both correlation and classification metrics (Figure 1C and D).

### Evaluation of shRNAI using publicly available shRNAmir libraries

A previous shRNAmir screening study to identify essential genes produced a large-scale dataset with a mir- E-like backbone (61) (Supplementary Figure 2A). Using an annotated essential gene set (76), both the shRNAI and SplashRNA models were evaluated by assessing how each model identified top and bottom shRNAmirs. This comparison showed that shRNAI had a better performance in detecting top-ranked shRNAmirs than SplashRNA, DSIR, and SeqScore (Figure 2A; one-sided Wilcoxon rank sum test). We then tested how accurately each model discerned true positives (essential genes) and true negatives (nonessential genes) using top-ranked shRNAmirs, and the results showed that shRNAI had a better ability to identify the top two shRNAmirs (Figure 2B).

**Figure 2.**
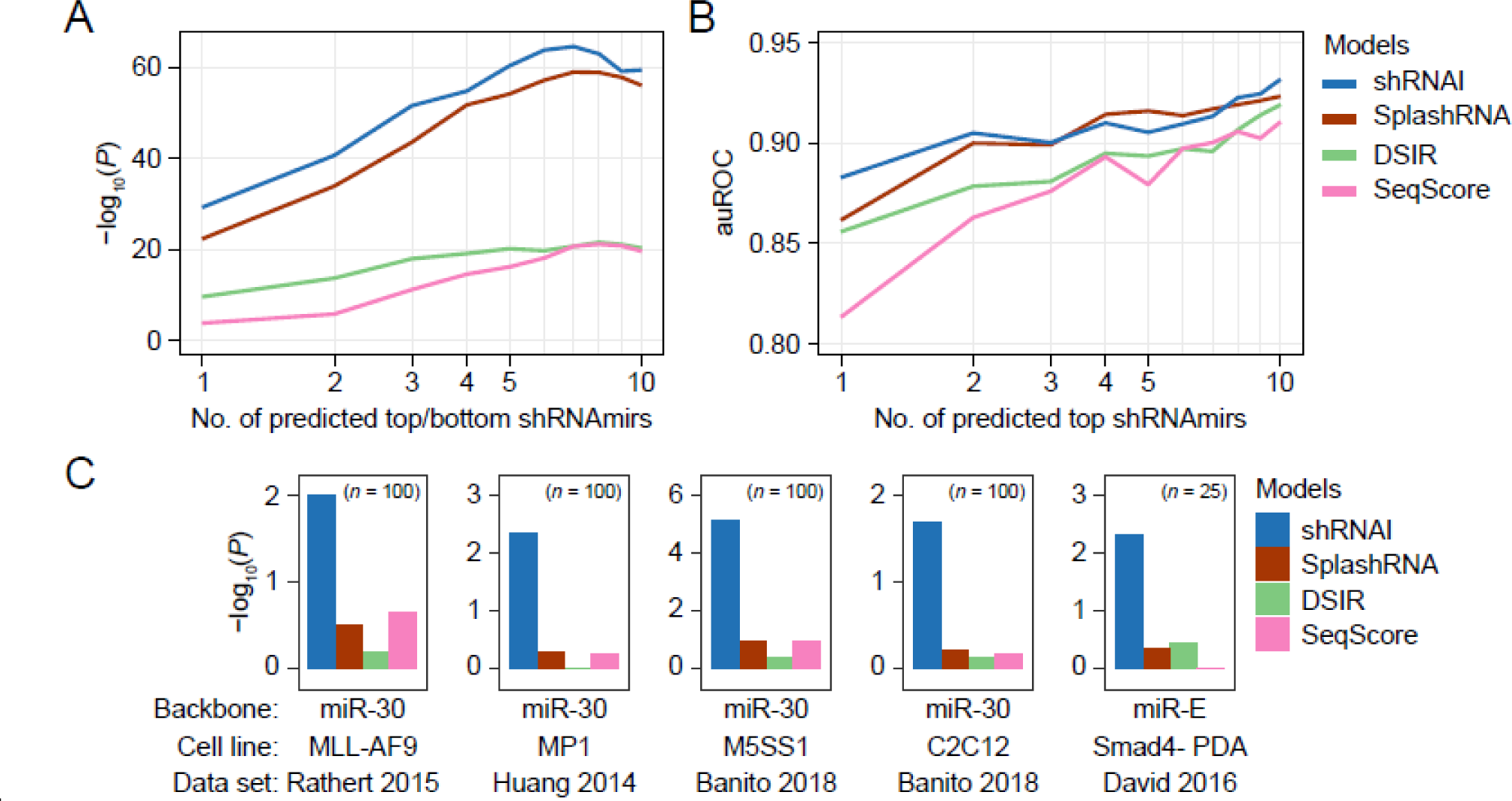
In silico validation of shRNAI performance. (A) Performance comparison of shRNAI with SplashRNA in retrospective shRNA potency prediction of essential gene sets. For each of the top 50 essential genes, all the tested algorithms selected top and bottom *N* sequences by prediction score and compared their real potency. *N* ranged from 1 to 10. Statistical significance was determined by a one-tailed Wilcoxon rank-sum test. (B) Performance comparison of shRNAI with SplashRNA in gold-standard essential and nonessential gene detection. The area under the receiver operating characteristic (auROC) curves were measured to detect hits in the top and bottom *N* sequences by the prediction score and compared. *N* ranged from 1 to 10. (C) Performance comparison of shRNAI with other algorithms in retrospective shRNA potency prediction of small-scale screening data. For each of the indicated genes, all the tested algorithms selected the top and bottom 1 sequences according to their prediction scores and compared their real potencies. Statistical significance was determined by a one-tailed Wilcoxon rank-sum test.

We further evaluated our shRNAI model with five small-scale screening datasets (Supplementary Figure 2B-O and Supplementary Table 2; see Methods for more details) (62–65). The observed potencies of shRNAmirs targeting 25 or 100 genes were compared to the best and worst potencies predicted by each benchmarking model. We found that shRNAI was the only model that significantly discerned the best and worst shRNAmirs across all the datasets (Figure 2C; one-sided Wilcoxon rank sum test), suggesting that the shRNAI model robustly outperformed the current methods.

### Prediction of shRNAmir processability by Drosha enhances the shRNAI model

Because shRNAmirs are processed by the endogenous miRNA processing pathway, we hypothesized that their potency could be affected by their processability, which includes the efficiency and accuracy of Drosha and Dicer processing. Hence, the relative abundances of primary shRNA (pri-shRNA, also referred to as shRNAmir), precursor shRNA (pre-shRNA), and mature gRNA were estimated from the TILE dataset. We conducted a correlation analysis between the relative abundance and potency of each shRNAmir (Supplementary Figure 3A-C). The relative levels of pri-shRNAs not processed into pre-shRNAs were negatively correlated with potency in the three stratified groups, while the relative levels of pre-shRNAs and gRNAs were not correlated. To identify the factors that cause stratified formation, the gRNAs were separated into three groups using two heuristic cutoff criteria (Supplementary Figure 3D). A comparison of the groups according to the position of the gRNAs revealed strong enrichment of AU content and 5’-end AU levels in the gRNAs with a high potency (Supplementary Figure 3E and F), as previously described (55).

Given that the transcription rate (*tr*) of shRNAmir plasmids is similar when the same promoter is used for the massive parallel assay, the abundance of shRNAmirs is likely to be solely affected by the Drosha processing rate (*pr*). Hence, we considered the relative level of pri-shRNAs as the surplus (*s* = *tr* – *pr*) of Drosha substrate (Figure 3A). In fact, the correlation between the AU content and the observed surplus, *s*, was reciprocal to that between the AU content and the Drosha cleavage score, as calculated from *in vitro* Drosha processing data (77) (Supplementary Figure 3G and H). Retraining the shRNAI model by adding *s* produced improved agreement with the observed potency in the TILE dataset (Figure 3B), indicating that the relative level of pri-shRNA is strongly correlated with Drosha cleavage efficiency. Given that the relative level of pri-shRNAs was only available in the TILE dataset and not in other datasets, we sought to predict the surplus value using another CNN model (Figure 3C; see Materials and Methods for more details). By considering the predicted surplus, *s*^, the shRNAI model was improved to a similar extent as when using the observed value (Figure 3D and E). The updated shRNAI***S*** model improved by approximately 9% compared to the SplashRNA model, which was retrained as described in the original paper (Supplementary Figure 3I), on the Ras dataset (Figure 3F; *r*=0.403 and *r*=0.370 for shRNAI and SplashRNA, respectively).

**Figure 3.**
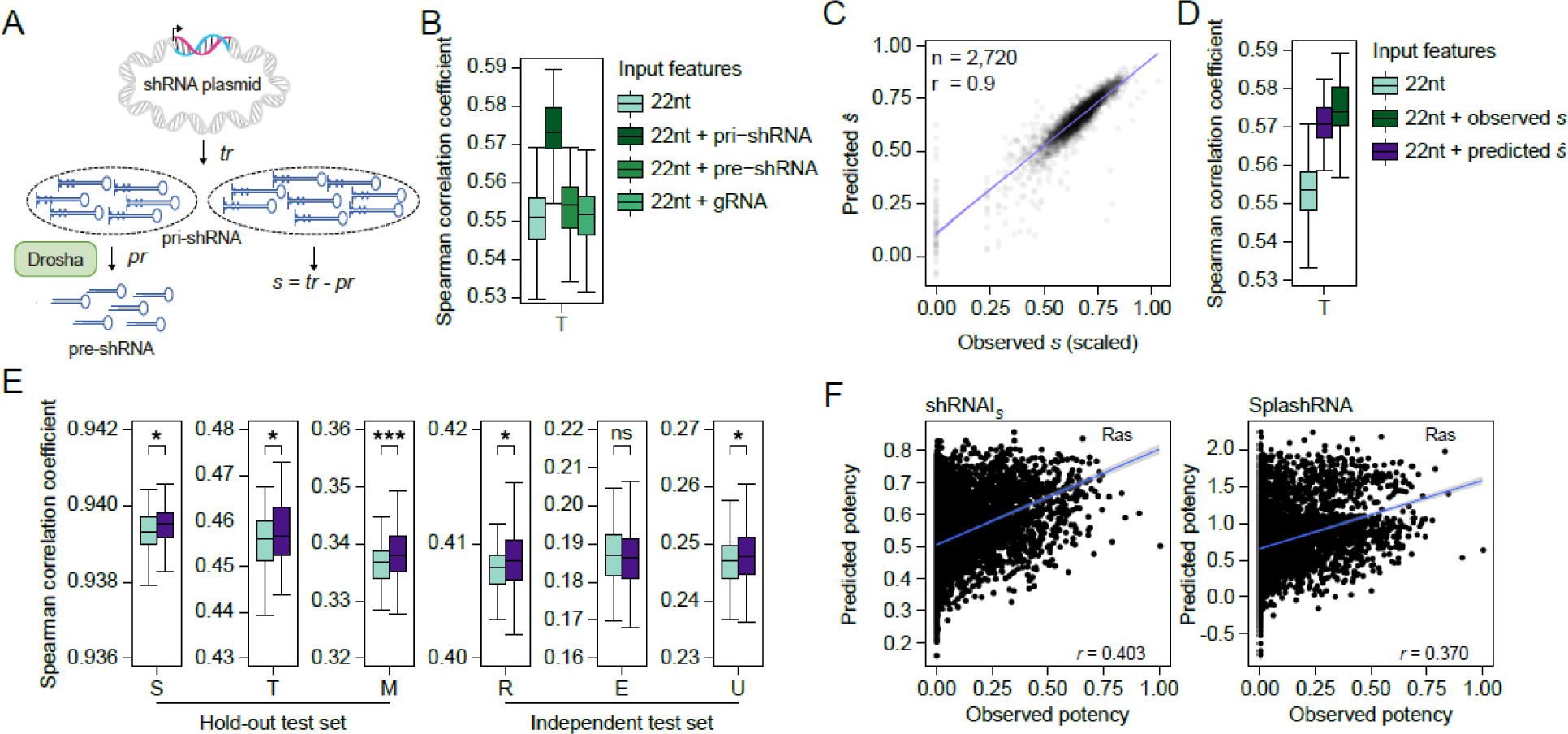
Improving model performance by considering Drosha processing efficiency. (A) Explanation of the Drosha substrate surplus (*s*). The variable *s* is calculated by subtracting the Drosha processing rate (*pr*) from the transcription rate (*tr*). (B) Performance comparisons of deep learning models trained on the training set of the T dataset and using its hold-out test set to test the contribution of additional features, including pri-shRNA, pre-shRNA, and gRNA abundance. (C) Scatter plot showing the correlation between the predicted and observed surpluses of Drosha substrates from the T hold-out test set. (D) Performance comparisons of deep learning models trained on the training set of the T dataset and using its hold-out test set to test the contributions of additional features, including predicted and observed surpluses of Drosha substrates. (E) Performance comparison between the sequence-based model and the model trained with the additional feature of the predicted surplus of Drosha substrate using a hold-out and an independent test set for all test datasets. Statistical significance was determined by a one-tailed Student’s t test. (F) Performance comparison between shRNAI*S* and SplashRNA on the Ras independent test set. Spearman correlation coefficients are indicated. The whiskers in the box plots in (B), (D), and (E) extend to 1.5x the interquartile range (the results are from 100 models trained with the same hyperparameters to show the convergence of the stochastic nature of building deep learning models).

### Experimental evaluation of shRNAI*S*

To experimentally evaluate shRNAI***S***, six target genes, *PTEN, BAP1, NF2, AXIN1, PBRM1*, and *RELA*, which were previously used for the evaluation of SplashRNA, were selected. For each target, five shRNAmirs with a higher shRNAI***S*** score than SplashRNA score (blue) and five shRNAmirs with a lower shRNAI***S*** score than SplashRNA score (red) were cloned and inserted into mir-E-based expression vectors and transduced into HeLa cells (multiplicity of infection (MOI) of 0.6) via a lentivirus system (Figure 4). We then assessed the knockdown (KD) efficiency using qRT‒PCR. The five shRNAmirs (blue) with a higher shRNAI***S*** score had a significantly greater KD efficiency than the five shRNAmirs (red) with a lower shRNAI***S*** score (*P* < 0.029 for *PTEN*; *P* < 0.017 for *BAP1*; *P* < 0.037 for *NF2*; *P* < 0.076 for *AXIN1*; *P* < 0.076 for *PBRM1*; *P* < 0.076 for *RELA*; one-sided Wilcoxon rank sum test); the five shRNAmirs (solid circle) with a higher SplashRNA score showed no significant differences with the five shRNAmirs (hollow triangle) with a lower SplashRNA score (*P* < 0.501 for *PTEN*; *P* < 0.891 for *BAP1*; *P* < 0.661 for *NF2*; *P* < 0.271 for *AXIN1*; *P* < 0.651 for *PBRM1*; *P* < 0.151 for *RELA*; Figure 4B, C; Supplementary Figure 4A- D; and Table S3). The shRNAI***S*** model exhibited a superior performance in all six targets according to the Spearman correlation between the prediction scores and observed values (Supplementary Figure 4E), suggesting that the shRNAI***S*** prediction results exhibited better agreement with the experimental results than did those of the SplashRNA model.

**Figure 4.**
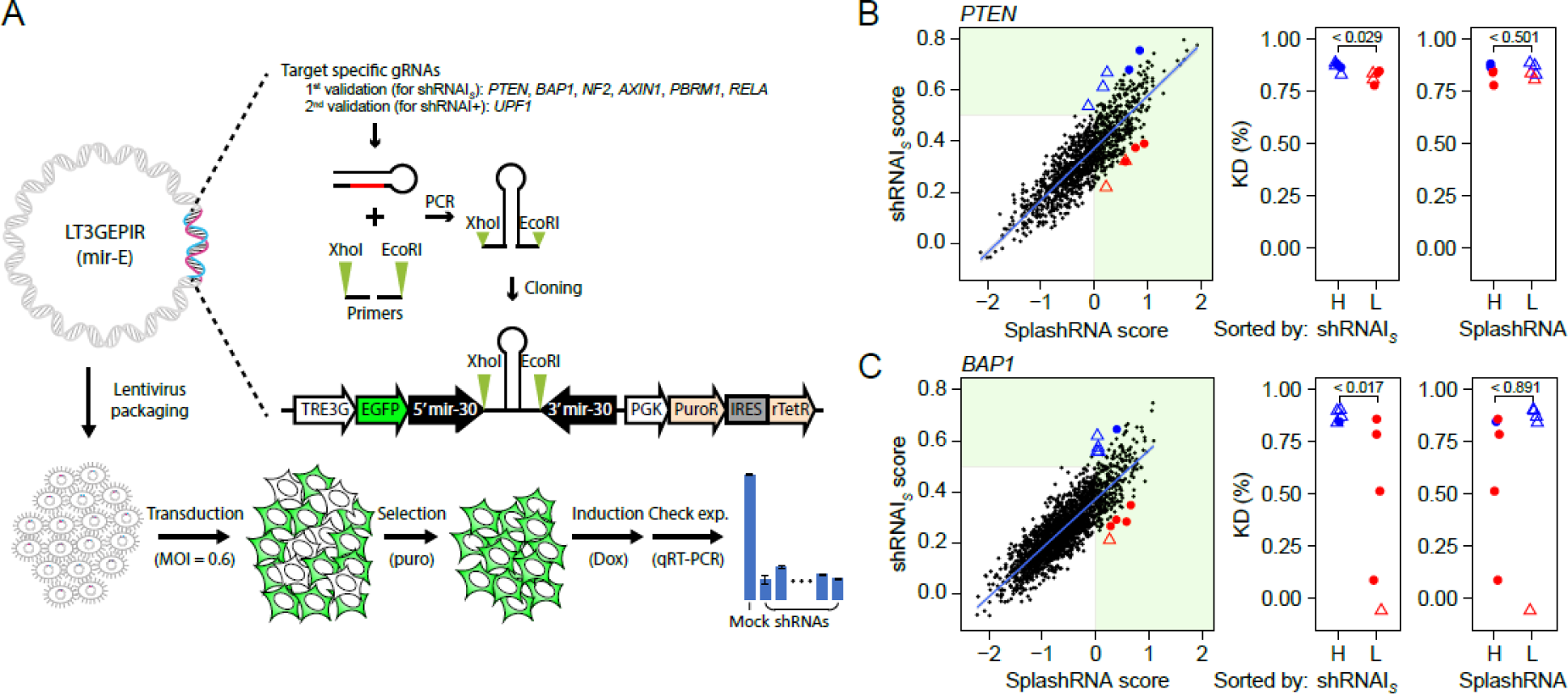
Experimental validation of shRNAI performance. (A) Illustration of the experimental validation process. The lentiviral and mir-E plasmids (LT3GEPIR) were utilized to evaluate the efficacy of the designed shRNAs. The gRNAs were selected based on criteria for a fair comparison between models and were cloned and inserted into the plasmid. The cloned plasmid was packaged into the lentivirus, and single-copy transduction was performed at an MOI of 0.6. After puromycin selection, qRT‒PCR of the target gene was performed to assess the knockdown efficiency. (B) Distribution of algorithm-predicted shRNA potency of the tiled CDS of *PTEN* (left). The x-axis indicates the SplashRNA score, and the y-axis indicates the shRNAI score. The selected shRNAmirs for validation are indicated as follows: blue indicates higher shRNAI scores and red indicates lower shRNAI scores; the solid circle indicates higher SplashRNA scores, and the hollow triangle indicates lower SplashRNA scores. Comparison of the knockdown (KD) efficiency between the two groups (right). Statistical significance was determined by a one-sided Wilcoxon rank-sum test. (C) Same as (B) but with shRNAmirs targeting *BAP1* instead of *PTEN*.

### Target site context improves the shRNAI model

The sequence contexts of the target sites and their flanking regions have been considered in detecting optimal target sites (78,79). Thus, we examined other datasets (S, T, M, and R) to determine whether the AU content in the flanking region of the target sites affects potency. Regardless of the dataset used, the target sites were consistently stratified by the number of A or U bases in the 2 nt downstream region, and the results showed that those with a greater AU content tended to have greater potency (Figure 5A). To provide comprehensive information on target site contexts to the model, the 14 nt sequence flanking the target site was incorporated as an input feature of the final shRNAI+ model. Regardless of the test dataset used, the shRNAI+ model incorporating the target site context outperformed the initial shRNAI model (Figure 5B). Further evaluation with independent test data from an earlier analysis (Figure 2) showed a comparable performance to the shRNAI model and a superior performance to SplashRNA (Figure 5C-E).

**Figure 5.**
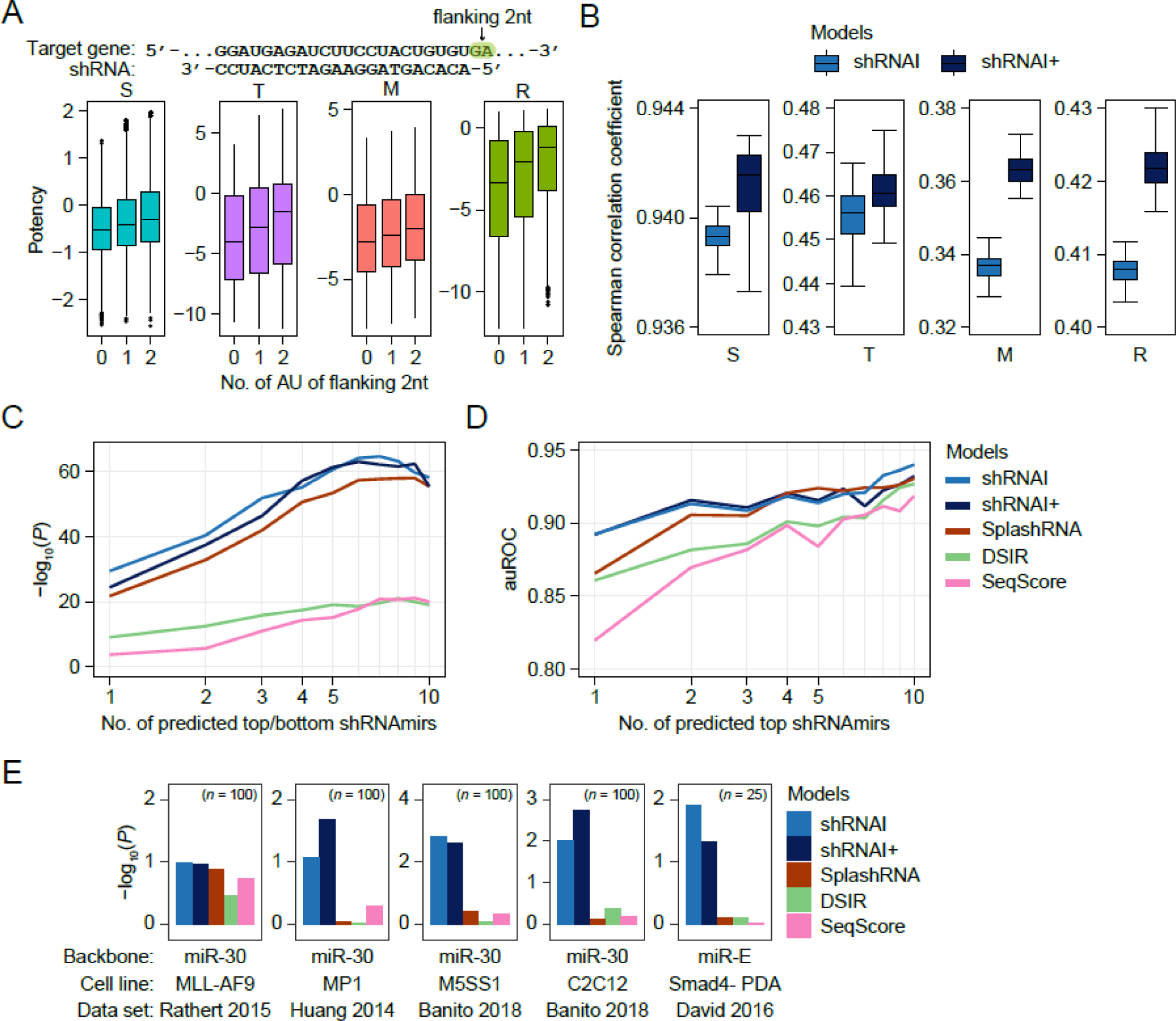
Improving model performance by considering the miRNA targeting mechanism. (A) (top) Illustration of the flanking 2 nt of the shRNA target site. (bottom) Comparison of shRNA potency distributions according to the number of AU nts in the downstream 2 nt region of the shRNA target site in each dataset. shRNAmirs with either A or U at the 5’ terminal of the mature siRNA and those with an AU content between 0.5 and 0.7 were used for this analysis. (B) Performance comparison between shRNAI and shRNAI+ using a hold-out and an independent test set for all datasets. (C) Same as Figure 2A but for the comparison of shRNAI+ with shRNAI and SplashRNA. (D) Same as Figure 2B but for the comparison of shRNAI+ with shRNAI and SplashRNA. (E) Same as Figure 2C but for the comparison of shRNAI+ with shRNAI and SplashRNA.

Next, after adding *UPF1* to our six target genes (Supplementary Figure 5A), a total of seven targets were used for experimental validation of the shRNAI+ model. All expression values of the target genes in the wild-type and shRNAmir treatment groups were normalized globally to the expression of *GAPDH*, followed by correlation with the predicted scores from the shRNAI+, shRNAI, and SplashRNA models. The shRNAI+ model predicted the experimental values better than the shRNAI and SplashRNA models (Figure 6A and B).

**Figure 6.**
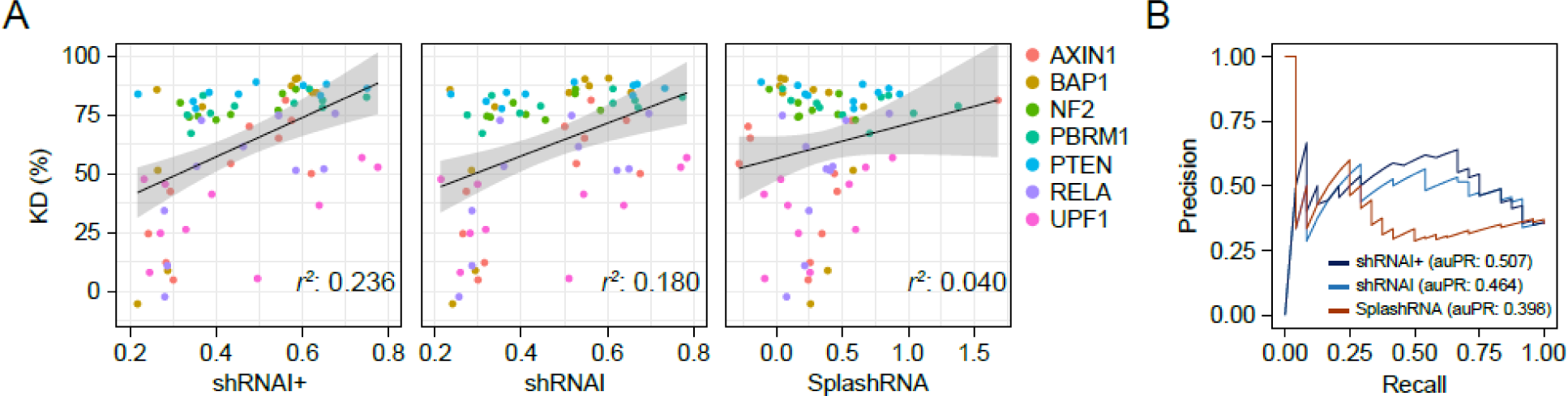
shRNAI+ performance test on experimental validation data. (A) The regression-based accuracies of KD efficiency were compared among three methods, shRNAI+ (left), shRNAI (middle), and SplashRNA (right), using shRNAmirs from Figure 4B and 4C and Supplementary Figures 4A-D and 6A. (B) Same as Figure 6A, but comparisons of classification-based accuracy were conducted instead of regression-based accuracy.

### The shRNAI+ model provides better siRNA-based drug candidates

Since the shRNAI+ model predicted the potency of gRNA loaded onto the shRNAmir backbone, we wanted to test whether this model could also be applied to predict better siRNAs using a large siRNA dataset (80). The correlation results with siRNA KD efficiency showed that our shRNAI models have predictive power for siRNA potency (Figure 7A). Since several siRNA-based drugs targeting *TTR*, *PCSK9*, *HAO1*, and *ALAS1* for the treatment of rare metabolic diseases were recently approved by the Food and Drug Administration (FDA) (81–85), we tried to identify potentially better gRNAs for these target genes using the shRNAI+ model. Our model detected better target sites and corresponding gRNAs with potentially greater potency than those of the FDA-approved siRNA-based drugs vutrisiran, patisiran, inclisiran, lumasiran, and givosiran (Figure 7B-E). To evaluate these target sites, we designed shRNAmirs against four target sites in the *TTR* gene that had shRNAI+ scores greater than 0.6 and two other sites targeted by FDA-approved siRNA drugs. The four shRNAmirs had a KD efficiency of more than 80%, and the best one had a KD of more than 90% in Huh7 cells, while the FDA-approved siRNA drugs had a KD efficiency of 84% and 70%, respectively (Figure 7F and Supplementary Table 4). These results suggest that the shRNAI+ model can be widely used for the development of siRNA- and shRNA-based drugs.

**Figure 7.**
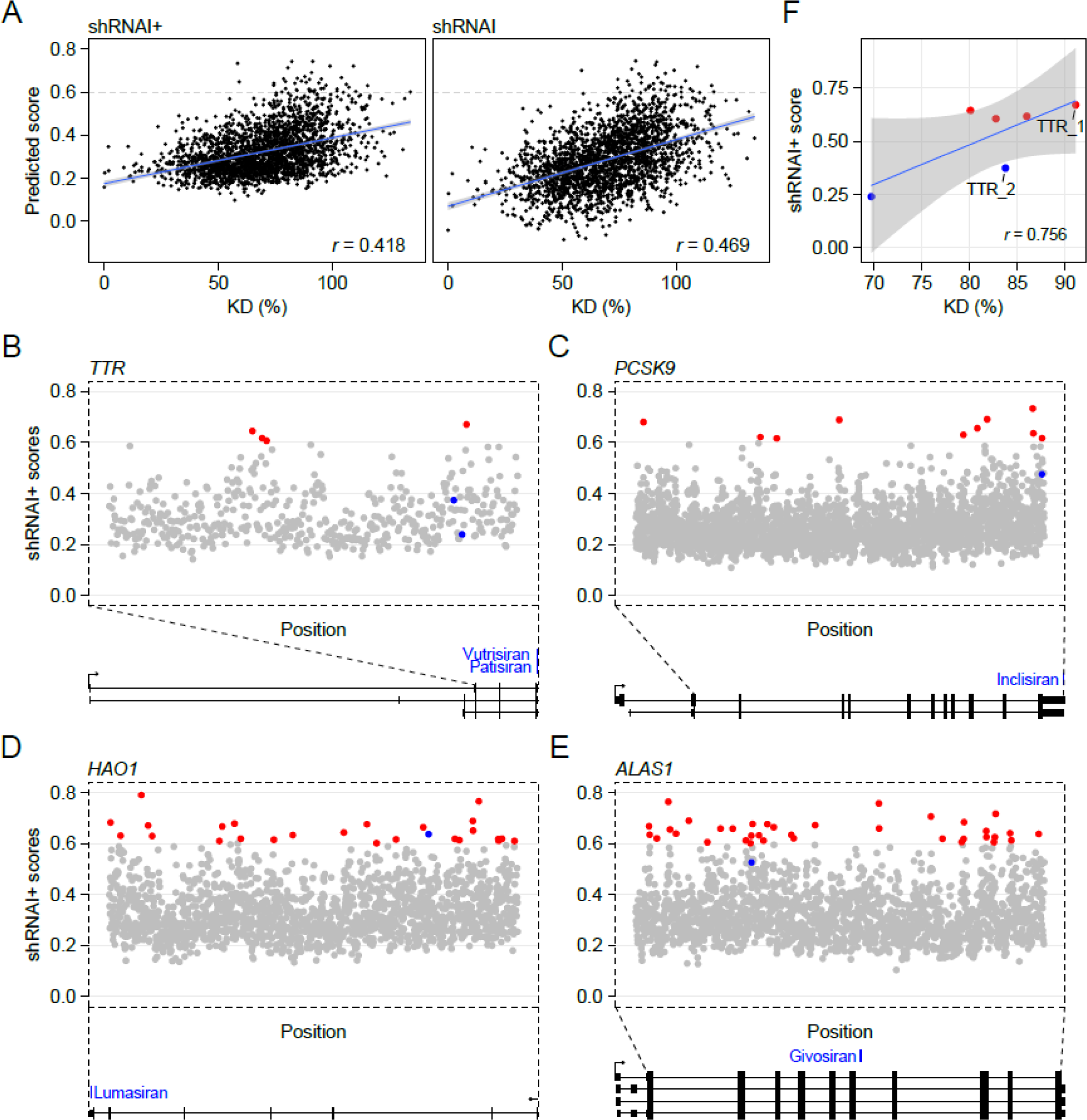
A suggestion for RNAi-based drug candidates using shRNAmirs. (A) Performance of shRNAI+ (left) and shRNAI (right) on siRNA screening data. Pearson correlation coefficients are indicated. (B-E) The bottom plots show the gene structures of *TTR* (B), *PCSK9* (C), *HAO1* (D), and *ALAS1* (E). The top plots show the distribution of the predicted shRNA potency of the tiled CDS and 3’UTR of each gene. The x-axis indicates the position, and the y-axis indicates the shRNAI+ score. The blue points indicate the same sites as those of FDA-approved siRNA drugs, and the red points indicate suggested target sites that showed high predicted potency. (F) Knockdown efficiencies of 6 selected shRNAmirs for the *TTR* gene, including 4 shRNAmirs with high shRNAI scores (red) and 2 shRNAmirs with the same target sites as FDA-approved siRNA drugs (blue). The Pearson correlation coefficient is indicated. Two shRNAmirs (TTR_1 and TTR_2) subjected to RNA- seq are also indicated.

To further investigate the potential off-target effects of shRNAmirs, we performed RNA-seq on cells transfected with our resulting shRNAmir (TTR_1) and the vutrisiran shRNAmir (TTR_2) (depicted in Figure 7F). We specifically examined the changes in the expression of mRNAs that could be off-targeted by shRNAmirs. These included gRNAs, isogRNAs (which are characterized by a 1 nt shift at the 5’ end of gRNA due to alternative processing), and pRNAs (passenger strands). We also considered the possibility of stoichiometric effects arising from competition with endogenous miRNAs, particularly focusing on the 2 most expressed endogenous miRNAs. We found minimal evidence of off-targeting and stoichiometric effects from the high-scoring shRNAmir (TTR_1), while the low-scoring shRNAmir (TTR_2) exhibited these effects with gRNAs, isogRNAs, and pRNAs (Supplementary Figure 7). These results suggest that well-designed shRNAmirs have promise as potent platforms for drug development, given their apparent specificity and minimal off-target effects.

### The CNNC motif may improve shRNAmir efficacy by enhancing Drosha processing

Although our models were trained with only mir-30 data lacking a CNNC motif, the shRNAI models still exhibited superior performance on public mir-E-based data (Figures 2 and 5) and our mir-E-based transduction data (Figures 4 and 6). To investigate how the CNNC motif influences shRNAmir potency, we reanalyzed publicly available *in vitro* Drosha processing data, which included approximately 100 billion variants in the 41–45 nt outside of the 8 nt upstream of the 5’ Drosha cleavage site and the 37–38 nt outside of the 8 nt downstream of the 3’ Drosha cleavage site of pri-miRNAs (pri-miR-125a, -miR-16-1, and -miR- 30) (86) (Supplementary Figure 6A). We first searched for sequence and structural motifs necessary for efficient pri-miRNA processing in the presence or absence of the CNNC motif. Comparing variant clusters with and without the CNNC motif based on the ratio of cleaved and input variants showed an additional effect on Drosha processing by pri-miR-30 and pri-miR-16, which had low mGHG scores (9), but not by pri-miR-125, which had a high mGHG score (95.93) (Supplementary Figure 6B). In particular, approximately half of the pri-miR-30 variants were found to be additively affected by the CNNC motif (sensitive vs. insensitive group; Supplementary Figure 7C), and the sensitive group displayed enriched motifs. Approximately 40% of the pri-miR-30 variant group sensitive to the CNNC motif included the base pair at position -12 from the 5’ Drosha cleavage site (termed pair12), while only approximately 5% of the variants insensitive to the CNNC motif had this base pair at this position (Supplementary Figure 6D, *FDR*<1.38x10^-18^; one-sided Fisher’s exact test). The pair12 motif exhibited enrichment of Watson–Crick base pairs A–T or G–C at the -12 position from the 5’ Drosha cleavage site in the CNNC motif-sensitive group (Supplementary Figure 6E, top). In contrast, T on the 3’ arm was significantly depleted at the -14 position from the 5’ Drosha cleavage site (Supplementary Figure 6E, bottom; *P*<1.1x10^-96^; one-sided binomial test), which was named the stem14 motif. Pri-miR-30 variant clusters with the pair12 motif or without the stem14 motif appeared to be sensitive to the CNNC motif (Supplementary Figure 6F). However, in the original paper (86), the authors insisted that the sensitivity of the CNNC motif to the presence or absence of pair12 and stem14 was not reproduced via *in vivo* expression. These results suggest that, regardless of the motif, the CNNC motif could enhance Drosha processing in the pri-miR-30 backbone, as shown for pri-miR-16 (Supplementary Figure 6B). In summary, the current shRNAI+ model does not consider the CNNC motif, which increases shRNAmir potency, due to the small mir-E dataset, but additional large amounts of data from mir-E backbone-based shRNAmirs could benefit the next version of the shRNAI+ model in the future.

## DISCUSSION

We implemented high-performance deep learning models, shRNAI and shRNAI+, to predict shRNAmir potency, and the models showed robustly superior performance in various independent test datasets. The improved model was generated by including shRNAmir processing efficiency as well as the context of the target sites and flanking regions. We found that a site with an shRNAI+ score ≥ 0.6 had a median KD efficiency ≥ 80% (Supplementary Figure 8A). For more than 80% of the human and mouse genes, the shRNAI+ model can predict at least one shRNAmir with a KD efficiency ≥ 80%, and for more than 60% of the genes, it can predict at least five shRNAmirs with a KD efficiency ≥ 80% (Supplementary Figure 8B), suggesting that our model is applicable to a majority of genes with high potency.

The abundance of pri-shRNAs is affected mostly by the production and Drosha processing rates. In the original experiments, however, shRNAmirs were transcribed with the same promoter in the expression vectors or virus; thus, their levels showed a linear relationship with shRNA potency (Supplementary Figure 3A). Within this framework, it is crucial to highlight that Drosha processing, which constitutes the initial phase of shRNAmir maturation, plays a pivotal role in determining the functionality of shRNAmirs. This underlines the significance of this step in the overall process. shRNA potency is directly affected by the abundance of gRNAs and is subsequently determined by compounding factors, such as pre-shRNA export, Dicer processing of pre-shRNAs, AGO loading of gRNAs, and target-directed miRNA decay (87–92). These findings suggest that there is still room for improvement by better mimicking *in vivo* Drosha/Dicer processing efficiency, pre-shRNA export and AGO loading efficiency. The elaborate design of a large-scale shRNAmir experiment in which the levels of shRNAmirs, pre-shRNAs, and shRNAs are simultaneously measured, as well as the level of the targets, could strengthen the power of the AI-driven shRNAI+ model in the future.

siRNAs have been explored as potential therapeutic agents, leading to the approval of several siRNA-based drugs by the FDA (81–85). Despite the advantages of shRNAmirs over siRNAs or shRNAs, the development of shRNAmir-based drugs has been hampered by the lack of efficient shRNAmir design tools. shRNAmirs have a superior ability to silence targets due to their vector-based, sustained expression and assimilation into the endogenous miRNA processing pathway (37–39). In addition, mass production of shRNAmirs is easier because they can be biologically synthesized in cells, whereas siRNAs require chemical synthesis and various modifications to achieve sufficient amounts and potency. Overall, the shRNAmir platform presents a promising opportunity for drug development, and our shRNAI models have the potential to make a significant contribution to the field.

## DATA AVAILABILITY

The raw sequencing data discussed in this publication have been deposited in the Korean Nucleotide Archive (KONA) under accession number KAP230669. All the Python and R codes used in this manuscript are available in the GitHub repository (https://github.com/ParkSJ-91/shRNAI).

## ACKNOWLEDGEMENTS

We thank all the Bioinformatics and Genomics (BIG) laboratory members for their helpful comments. *Author contributions*. SJP performed the computational analysis, and SHP performed the ex vivo validation and analysis. SJP contributed to the writing of the codes. SJP, SHP, JKH, and JWN designed this study and contributed to the writing of the manuscript. JWM and JKH supervised the project. JWN and SJP conceived the idea.

## FUNDING

This work was supported by the Bio & Medical Technology Development Program and Basic Research Program of the National Research Foundation (NRF) funded by the Ministry of Science and ICT (Grant numbers 2021M3A9I4024452 and 2020R1I1A2075393 to J.K.H. and 2020R1A4A1018398, 2021R1A2C3005835, 2022M3E5F1018502, RS-2023-00207840, and 2023R1A6C101A009 to J.W.N.).

This research was also supported by a grant from the Korea Health Technology R&D Project through the Korea Health Industry Development Institute (KHIDI), funded by the Ministry of Health & Welfare, Republic of Korea (grant number: HI22C063600 to J.K.H.).

## Conflict of interest statement

The authors declare no competing interests.

## Supplementary Information

**Figure S1.**
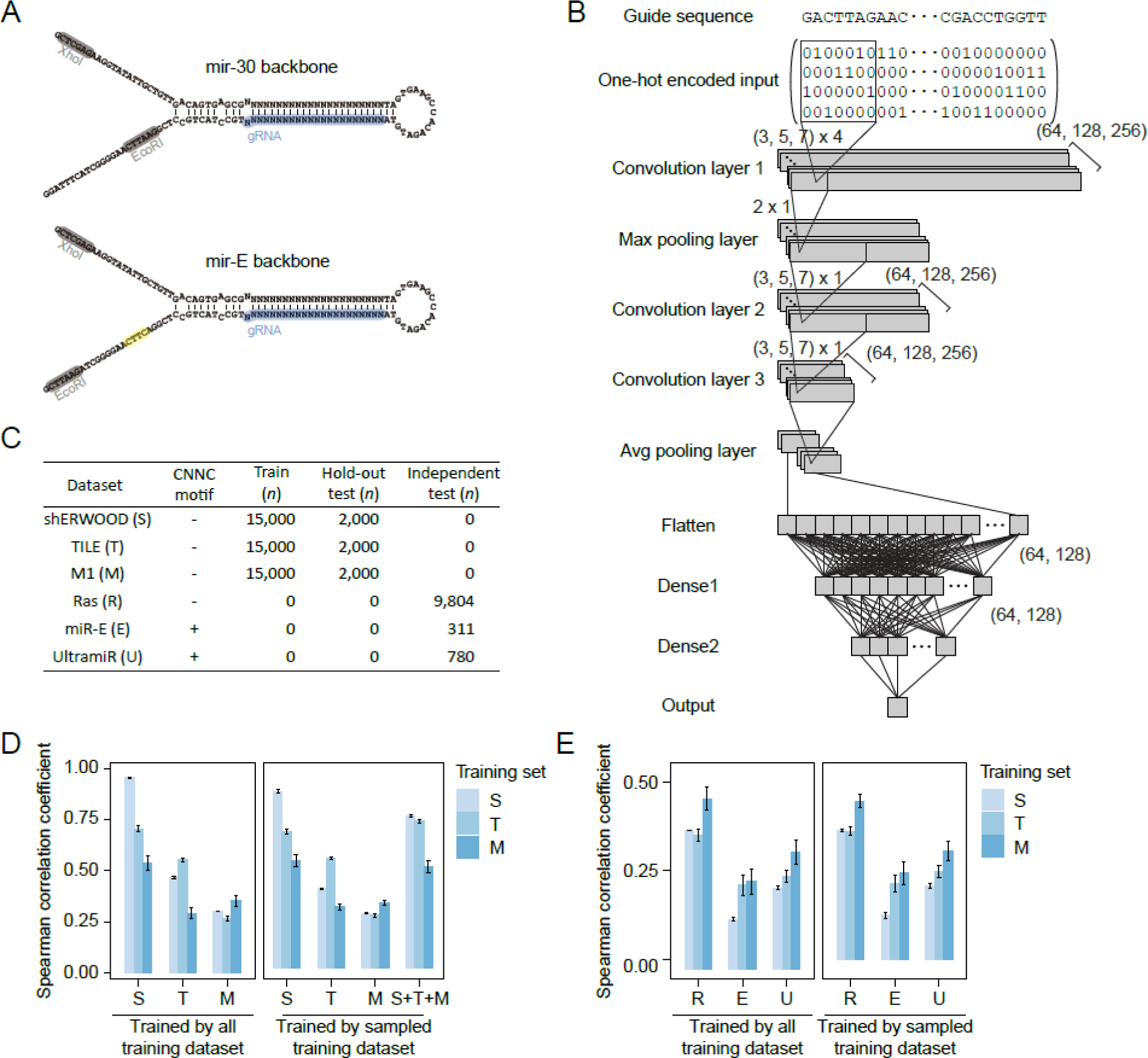
**Implementing CNN-based shRNAI.** (A) The sequence and structure of the mir-30 and mir-E backbones. (B) Topology of the convolutional neural network for implementing shRNAI consisting of three convolutional layers interlaced with a max pooling layer, two fully connected layers, and a regression output node. shRNAI uses one-hot encoded 22-nt guide sequences as input to predict the potency of each shRNA. (C) Performance comparison of deep learning models trained on the full training set (left) or the sampled training set (right) using the hold-out test set. The data represent the median ± SD for 100 models trained with the same hyperparameters to show the convergence of the stochastic nature of building deep learning models. (D) The number of training and testing data points sampled to select the appropriate combination. Training and hold-out test data were randomly sampled from full data points in the training set and hold-out test set, respectively, in the shERWOOD, TILE, and M1 datasets (Figure 1B). For the independent test set, all the data points in the Ras, miR-E, and UltramiR datasets were used. (E) Performance comparison of deep learning models trained on the full training set (left) or the sampled training set (right) using an independent test set. The data represent the median ± SD for 100 models trained with the same hyperparameters to show the convergence of the stochastic nature of building deep learning models.

**Figure S2.**
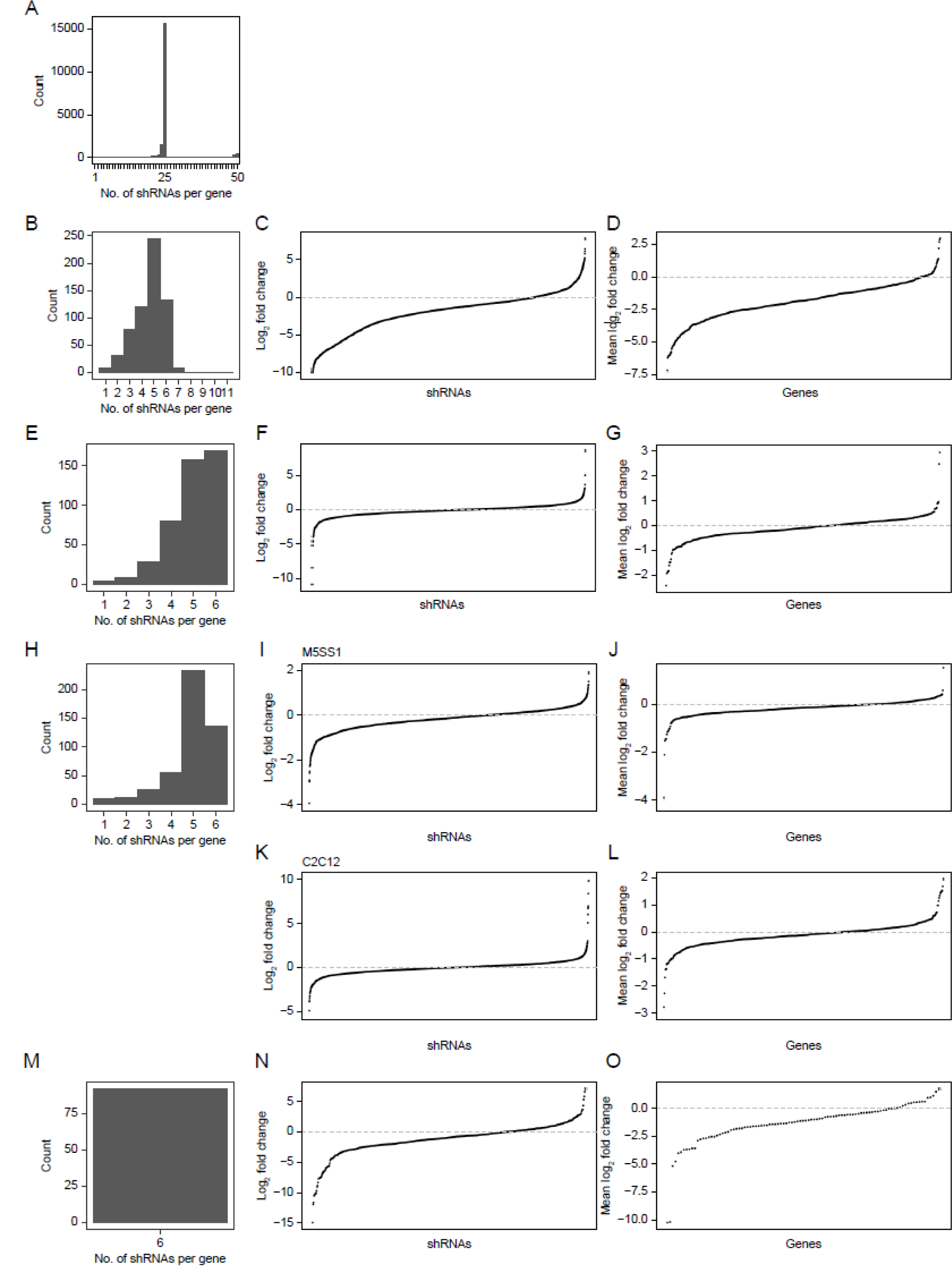
Distribution of the screening dataset. (A) Distribution of the number of shRNAs per gene in the essential gene set. (B) Distribution of the number of shRNAs per gene in the Rathert 2015 cohort. (C) Differences in shRNA representation are presented as log2 values of the ratio between average reads of the sequencing data replicates generated before and after treatment with shRNAs in the Rathert 2015 set. (D) Differences in gene representation are presented as the average shRNA representation in Supplementary Figure 2C. (E) Distribution of the number of shRNAs per gene in the Huang 2014 cohort. (F) Differences in shRNA representation are presented as log2 values of the ratio between average reads of the sequencing data replicates generated before and after treatment with shRNAs in the Huang 2014 set. (G) Differences in gene representation are presented as the average shRNA representation in Supplementary Figure 2F. (H) Distribution of the number of shRNAs per gene in the Banito 2018 cohort. (I) Differences in shRNA representation are presented as log2 values of the ratio between average reads of the sequencing data replicates generated before and after treatment with shRNAs in M5SS1 cells in the Banito 2018 cohort. (J) Differences in gene representation are presented as the average shRNA representation in Supplementary Figure 2I. (K) Differences in shRNA representation are presented as log2 values of the ratio between average reads of the sequencing data replicates generated before and after treatment with shRNAs in C2C12 cells in the Banito 2018 cohort. (L) Differences in gene representation are presented as the average shRNA representation in Supplementary Figure 2K. (M) Distribution of the number of shRNAs per gene in the David 2016 cohort. (N) Differences in shRNA representation are presented as log2 values of the ratio between average reads of the sequencing data replicates generated before and after treatment with shRNAs in the David 2016 cohort. (O) Differences in gene representation are presented as the average shRNA representation in Supplementary Figure 2M.

**Figure S3.**
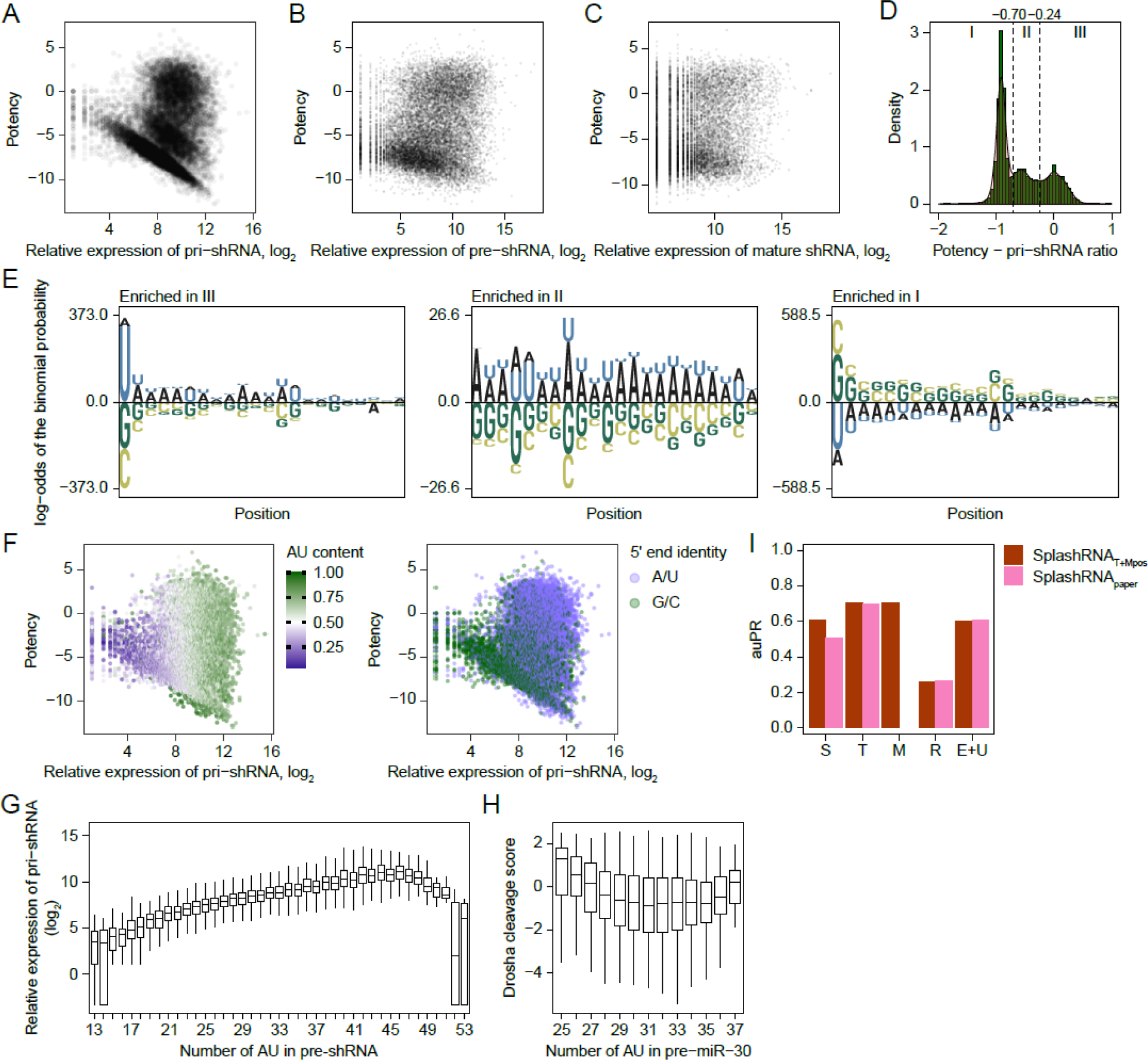
Stratified correlation between pri-shRNA abundance and potency. (A-C) Scatter plot showing the correlation between potency and the relative abundance of pri-shRNA (A), pre-shRNA (B), or mature gRNA (C). (D) Distribution of the ratio of the potency to the relative abundance of pri-shRNA. Heuristic cutoffs were used to separate the clusters (-070 and -0.24). (E) pLogo (93) results for enriched sequence compositions between clusters according to the position of the guide sequence. (F) Same plot as in Supplementary Figure 3A, but each guide sequence is colored according to the AU content (left) and 5’ end identity (right). (G) Correlation between the number of AU pairs in guide RNA and the relative abundance of pri-shRNA in the T dataset. (H) Correlation between the number of AU pairs in pre-miR-30 and the Drosha cleavage score determined from *in vitro* Drosha processing data. (I) Performance comparison between retrained SplashRNA (SplashRNAT+Mpos) and the original SplashRNApaper.

**Figure S4.**
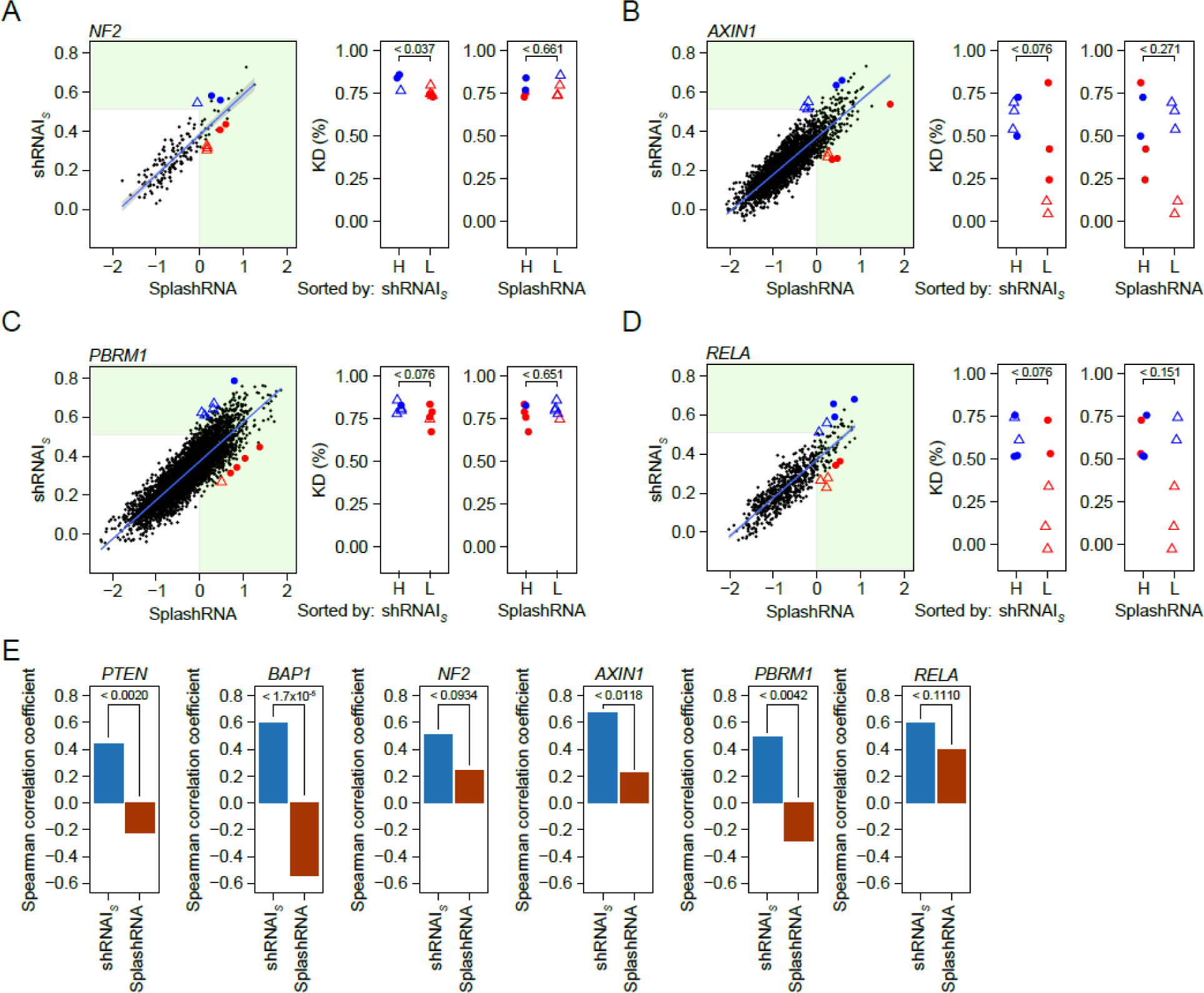
Experimental validation of shRNAI performance. (A-D) Same as Figure 4B, including shRNAmirs for *NF2* (A), *AXIN1* (B), *PBRM1* (C), and *RELA* (D) instead of *PTEN*. (E) Comparison of Spearman correlation coefficients between knockdown (KD) efficiency and the scores predicted by each model. Statistical significance was determined by a one-sided Steiger’s test.

**Figure S5.**
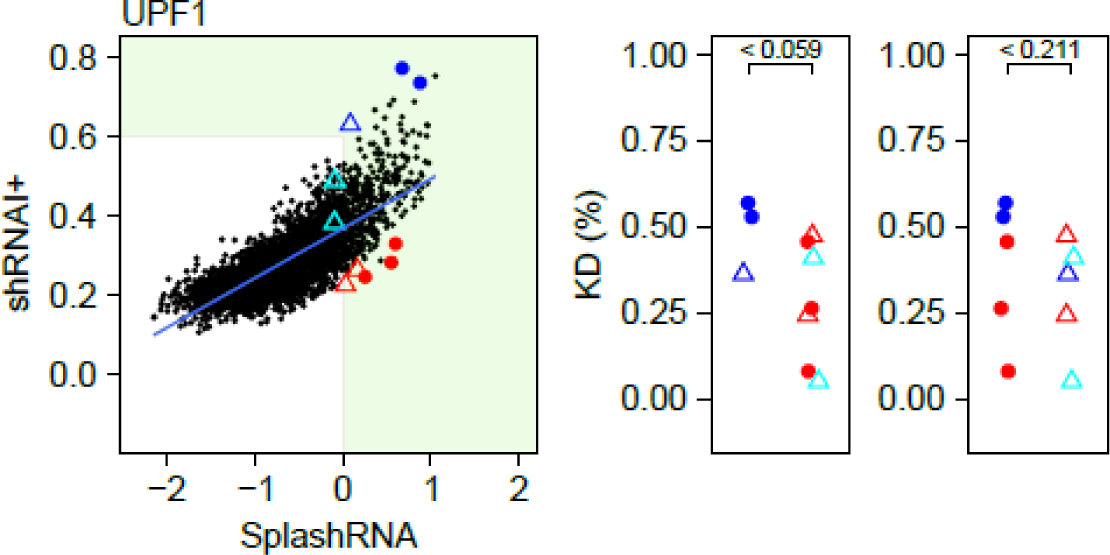
Analysis of the experimental validation results to estimate the cutoff using bootstrapping. Same as Figure 4B but with shRNAmirs for *UPF1* instead of *PTEN* and shRNAI+ instead of shRNAI.

**Figure S6.**
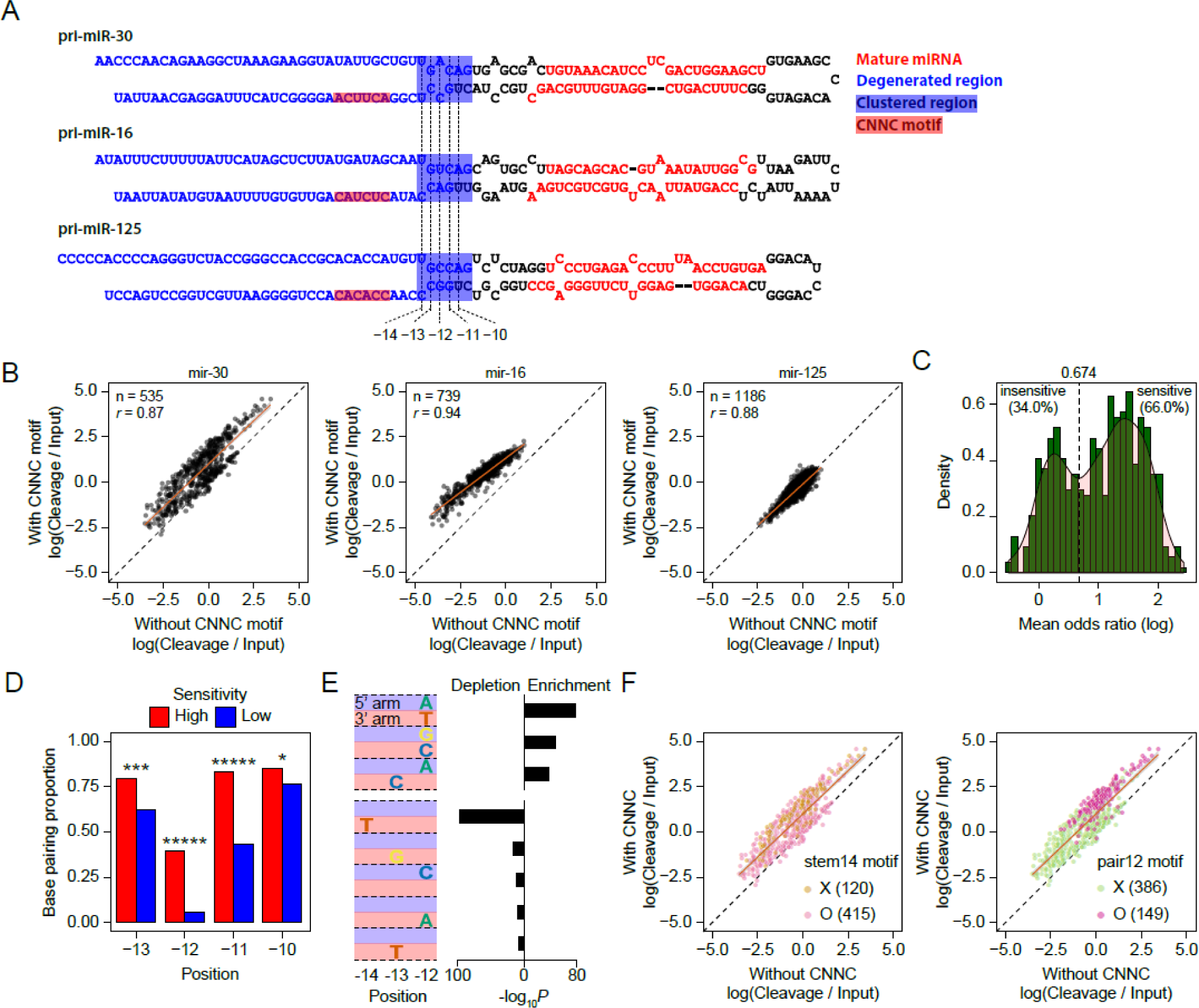
Evidence that the effect of the CNNC motif on Drosha processing efficiency is independent of the gRNA sequence. (A) The sequences and secondary structures of the three pri-miRNAs. The regions of both mature miRNAs are colored red, and the degenerate sequences are colored blue. The clustered region and the CNNC motif region are indicated by squares and colored blue and red, respectively. (B) Scatter plots showing the correlation of the cleavage ratio over the initial population (input) between clusters with and without the CNNC motif. (C) Distribution of odds ratios for cleavage ratios over the initial population within clusters with the CNNC motif relative to those without the CNNC motif. A heuristic cutoff (0.674) was used to separate clusters into insensitive and sensitive groups. (D) Comparison of mismatch proportions between the CNNC motif-sensitive and insensitive groups at each position in the clustered region. Statistical significance was determined by Fisher’s exact test. *: P ≤ 0.05, ***: P ≤ 0.001, ****: P ≤ 0.0001, and *****: P ≤ 0.00001. (E) *kp*Logo (94) results for enriched 2-mer (top) and depleted 1-mer (bottom) sequence compositions between the CNNC motif-sensitive and insensitive clusters according to the position of the clustered region. (F) Same plot as in Supplementary Figure 6B (left) but colored according to the presence of the stem14 motif (left) or the pair12 motif (right).

**Figure S7.**
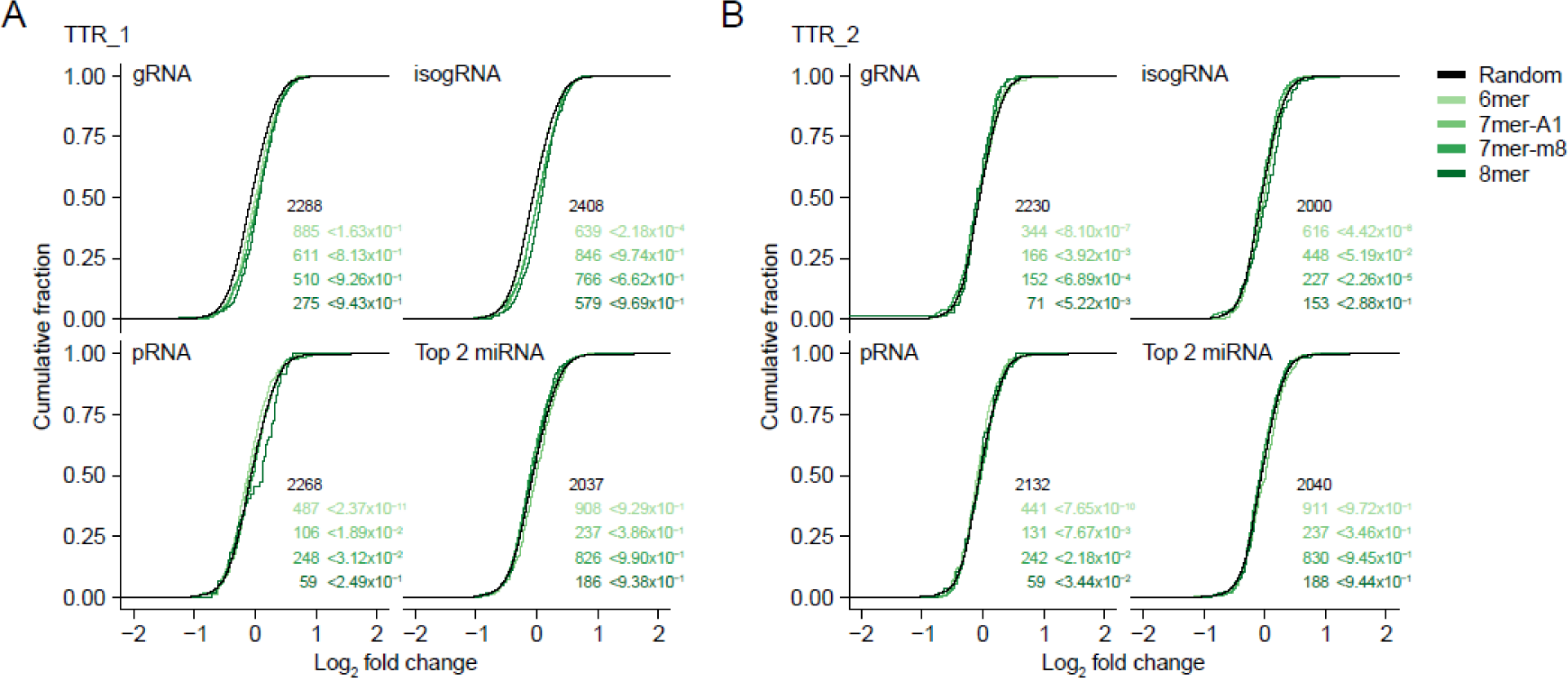
Minimal off-target effects of shRNAmirs. Response of mRNAs to the transfection of shRNAmirs TTR_1 (A) and TTR_2 (B), comparing mRNAs that contain a canonical miRNA target site to AGO-loadable RNAs, including gRNA, isogRNA, pRNA, and the top 2 endogenous miRNAs, to those that contain a canonical miRNA target site of 10 dinucleotide shuffled sequences. Cumulative distributions of mRNA fold changes calculated by comparing cells transfected with negative control shRNA to those transfected with TTR-targeting shRNAs are plotted. A one-sided Kolmogorov‒Smirnov test was conducted between site-containing distributions and random site distributions. The number of mRNAs analyzed in each category is listed in the graph.

**Figure S8.**
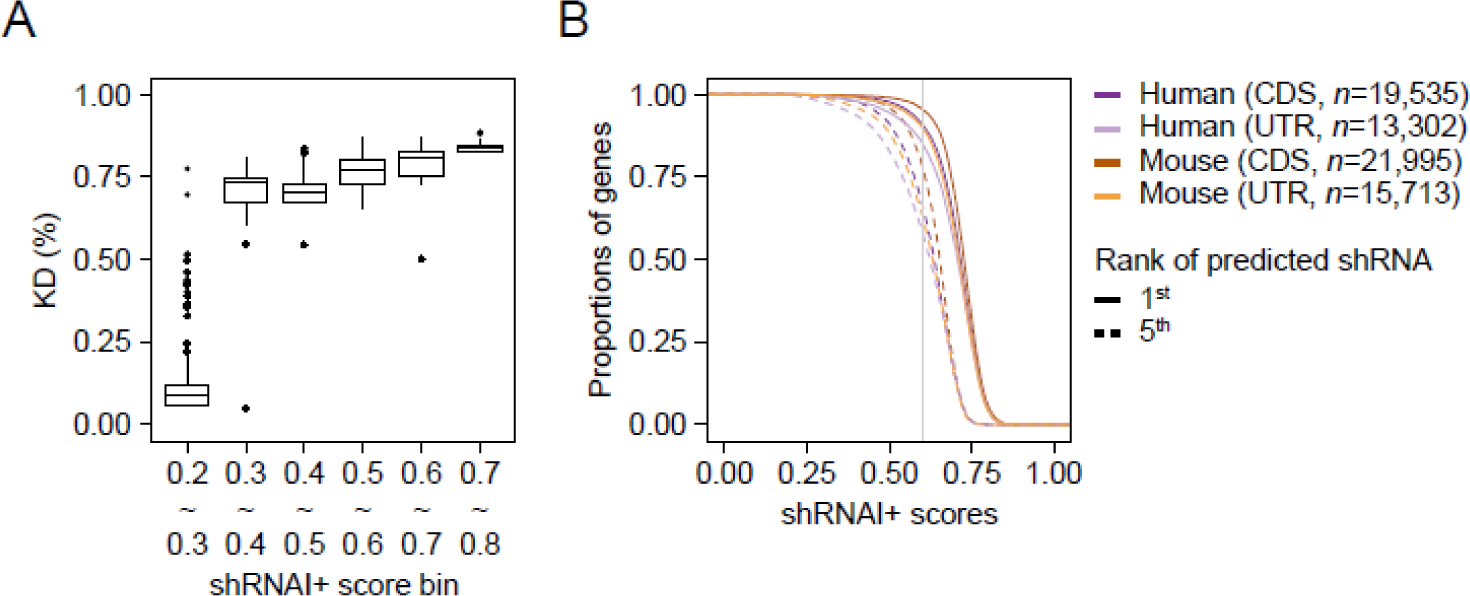
Cutoff of the shRNAI+ score for potent gRNA design. (A) The distribution of 80th percentile values from 10,000 bootstrapping iterations on the experimental validation results. In each iteration, 10 data points were resampled across 6 bins from the original data. The x-axis represents the bin, and the y-axis represents the 80th percentile value. (B) Score distribution of the first highest or fifth highest shRNAI predictions for all human and mouse genes.

**Table S1. Existing potency datasets used for training and testing shRNAI.** The number of total data points in each dataset is shown along with the number of positive and negative points divided by the indicated thresholds used in SplashRNA. The same thresholds were used to test the classification performance of shRNAI.

**Table S2. Small-scale screening datasets.** The number of total target genes and shRNAs used in each dataset are shown, along with a brief description of the experiment, including the cell lines used and the selection type.

**Table S3. Experimental validation results.** The KD efficiencies of the tested shRNAs are shown in triplicate.

**Table S4. Drug candidate screening results.** The KD efficiencies of the tested shRNAs are shown in triplicate.

## Notes

### Competing Interest Statement

The authors have declared no competing interest.

